# Transcriptome-wide high-throughput mapping of protein-RNA occupancy profiles using POP-seq

**DOI:** 10.1101/2020.12.28.424570

**Authors:** Mansi Srivastava, Rajneesh Srivastava, Sarath Chandra Janga

## Abstract

Interaction between proteins and RNA is critical for post-transcriptional regulatory processes. Existing high throughput methods based on crosslinking of the protein-RNA complexes and polyA pull down are reported to contribute to biases and are not readily amenable for identifying interaction sites on non polyA RNAs. We present Protein Occupancy Profile-Sequencing (POP-seq), a phase separation based method in three versions, one of which does not require crosslinking, thus providing unbiased protein occupancy profiles on whole cell transcriptome without the requirement of polyA pulldown. Our study demonstrates that ~68% of the total POP-seq peaks exhibited an overlap with publicly available protein-RNA interaction profiles of 97 RNA binding proteins (RBPs) in K562 cells. We show that POP-seq variants consistently capture protein-RNA interaction sites across a broad range of genes including on transcripts encoding for transcription factors (TFs), RNA-Binding Proteins (RBPs) and long non-coding RNAs (lncRNAs). POP-seq identified peaks exhibited a significant enrichment (p value < 2.2e-16) for GWAS SNPs, phenotypic, clinically relevant germline as well as somatic variants reported in cancer genomes, suggesting the prevalence of uncharacterized genomic variation in protein occupied sites on RNA. We demonstrate that the abundance of POP-seq peaks increases with an increase in expression of lncRNAs, suggesting that highly expressed lncRNA are likely to act as sponges for RBPs, contributing to the rewiring of protein-RNA interaction network in cancer cells. Overall, our data supports POP-seq as a robust and cost-effective method that could be applied to primary tissues for mapping global protein occupancies.

## Introduction

Interaction of proteins with RNA is crucial for post-transcriptional gene regulation such as capping, splicing, polyadenylation and localization which is indispensable for cellular homeostasis and survival^1,2^. Despite the increasingly appreciated role of protein-RNA interactions, the global occupancy profiles of proteins in a cellular environment is not fully elucidated. For instance dysregulated expression of RNA binding proteins (RBPs) has been associated with a broad spectrum of human pathologies including cancers, neurological and hereditary diseases^3–6^. Therefore, it is critical to investigate the diverse protein occupancy sites and their functional impact on physiology and diseases.

Experimental approaches such as crosslinking followed by immunoprecipitation (CLIP) have been widely used to identify the binding pockets of specific RBPs across the transcriptome^7,8^. CLIP based methods exploit the stability of crosslinked protein-RNA complexes by ultraviolet (UV) irradiation followed by immunoprecipitation and sequencing of the co-purified RNA^9–11^. However, these methods are reported to contribute to biases in the interaction profiles due to the inherent nature of UV crosslinking^12–14^ Other methods that employ antibody pulldown of protein-RNA complexes such as RIP-seq and DO-RIP-seq^15–17^ are difficult to scale up for detecting hundreds of RNA interacting proteins at the same time. Methods like POPPI-seq^18^ and others have enabled the capture of protein-RNA interaction sites by incorporation of photoreactive nucleosides in UV irradiated cells followed by poly-A pull down and sequencing of the bound RNA. Although, the use of photoreactive nucleosides is known to induce cellular stress that may result in non-physiological protein-RNA interactions and thus limits their application to only in-vitro cultures^8^. In addition, polyA pulldown requirement in these methods excludes their application to non-polyadenylated RNAs such as lncRNAs, miRNAs and histone mRNAs^18,19^. Several methods also employ formaldehyde crosslinking to capture protein-RNA complexes, however formaldehyde is also known to introduce biases by capturing non-specific interactions^8, 20–22^. Therefore, there is a need to develop unbiased and cost-effective method that can map the global occupancy profiles of protein bound sites in a transcriptome wide manner with its application to diverse species of RNA.

Trizol based phase separation strategy has emerged as a robust technology that has expanded the identification of protein binding sites independent of the poly-A capture^23–25^. Trizol extraction is deployed as a prevalent method to purify total RNA from the cell lysates. This involves the solubilization of biological material by phenol and guanidium isothiocyanate followed by chloroform induced phase separation. After the phase separation, proteins migrate to the organic phase, RNA migrates to the aqueous phase and the DNA/RNA-protein adducts are trapped in the interphase.

In this study, we propose a method called POP-seq (Protein Occupancy Profile-sequencing) that incorporates a multi-step phase separation strategy using trizol, followed by high-throughput sequencing of small RNAs, without the requirement of poly-A pulldown. Current study reports three versions of POP-seq; NPOP-seq (no-crosslinking), FPOP-seq (formaldehyde crosslinking) and UPOP-seq (UV crosslinking), among which NPOP-seq can efficiently capture the interactions without crosslinking mediated biases under physiological conditions. Computational analysis of POP-seq data revealed a significant enrichment of clinically relevant somatic variants in the protein-RNA interaction sites. Further, this study also demonstrates that highly expressed lncRNAs act as sponges to titrate the abundance of RBPs thereby altering the protein-RNA regulatory networks. Overall, POP-seq is a robust and cost-effective method that can be utilized by researchers to capture the protein occupied sites on all RNA types.

## Results

### POP-seq captures protein bound RNA fragments with three transcriptome wide approaches

We aimed to generate an unbiased transcriptome wide protein-RNA occupancy profiles using a trizol based phase separation method, POP-seq (Protein Occupancy Profile-sequencing) in K562 cells. Recent studies have demonstrated that phase separation using trizol yields abundance of RNA-protein interactions at the interphase^23,24^ However, identification of precise protein occupied pockets across transcriptome remains obscure.

POP-seq is employed in three different versions: NPOP-seq (no crosslinking), FPOP-seq (Formaldehyde crosslinking) and UPOP-seq (UV crosslinking) in K562 cells (Figure 1A). POP-seq employs trizol lysis of cells that generates three phases: aqueous phase, interphase, and organic phase. After removal of aqueous and organic phases, interphase is subjected to RNase A/T1 digestion, to remove the unprotected RNA from the RNA-Protein complexes trapped in the interphase. This is followed by degradation of the bound protein counterpart leaving behind the small RNA pockets using proteinase K. Further, DNase treatment ensures the sample quality by eliminating any DNA traces that might arise from the interphase. This is followed by removal of highly abundant ribosomal RNA (Figure 1B). Implementation of RNase digestion creates a 5’-hydroxyl, and 3’-phosphate ends in purified RNA making it inappropriate for adapter ligation during library preparation. Therefore, we modified RNA ends using T4 Polynucleotide kinase (PNK) and Calf intestine alkaline phosphatase (CIAP) to add 5’-phosphate and 3’-hydroxyl to the ends (Figure 1B). RNA integrity was assessed by Bioanalyzer QC and small RNA libraries were prepared for high throughput short read sequencing by Illumina, next-seq platform (Figure 1B). Together, this method allows identification of global protein occupancy across transcriptome.

**Figure 1.**
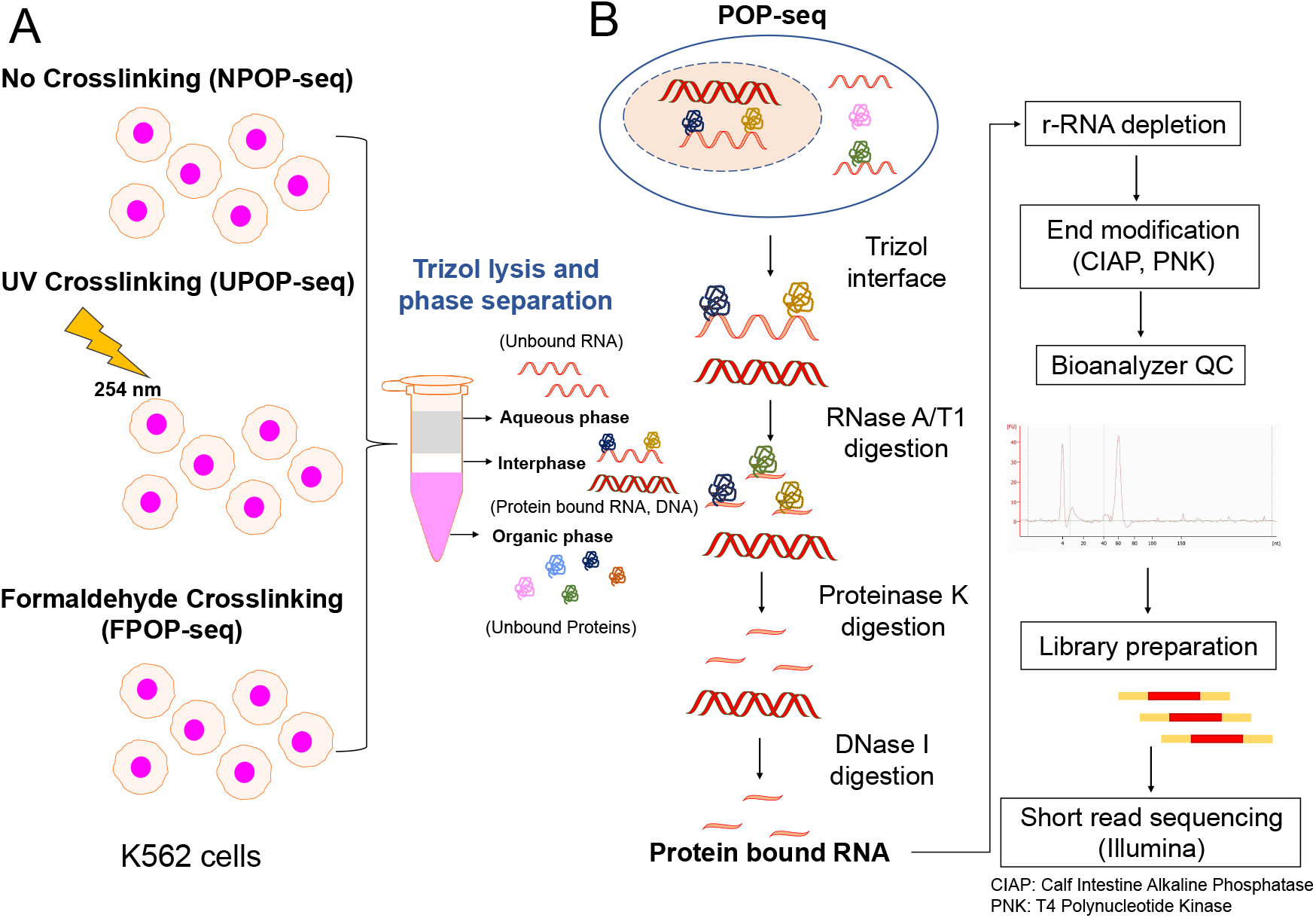
Experimental workflow of POP-seq in K562 cells. (A) Three versions of POP-seq, No-crosslinking (NPOP-seq), UV crosslinking, 254 nm (UPOP-seq) and formaldehyde crosslinking (FPOP-seq) generate three phases (aqueous, interphase and organic phase) upon trizol lysis, (B) Cell lysates from the POP-seq are digested with RNase A/T1 mix, Proteinase K and DNase followed by r-RNA depletion, RNA quality check and library preparation for short read Illumina sequencing.

### POP-seq reproducibly capture the abundance of protein-RNA interactions on the exonic regions in a human leukemia cell line

To characterize the transcriptome wide binding of the proteins, POP-seq libraries (in replicates) were sequenced to generate ~20 million reads each. We implemented our NGS pipeline to facilitate the analysis of the POP-seq data which includes quality control and read alignment followed by peak calling, resulting in the identification of 319657, 288129, and 320310 unique peaks in the respective protocols (Figure 2A). Overall, ~85% of total peaks had length below 50 bp (Figure 2B). Since reproducibility is an important aspect to estimate the robustness of high throughput methods, we compared the aligned reads per 10 kb genome in replicates for each protocol using deepTools^26^ ‘plot correlation’ command. Our data showed a correlation of ~64% (spearman correlation R^2^ values; 0.65, 0.64, 0.64 for NPOP, FPOP and UPOP respectively) between the replicates as shown in Figure S1. Comparison of POP-seq peaks with an end to end 50% peak overlap the combined eCLIP^27^ profile of 97 RBPs from the ENCODE project^28^ revealed support for 68.2%, 67.3% and 66.4% in NPOP, FPOP and UPOP-seq respectively. Additionally, we observed that ~64% of the genes exhibited by POP-seq peaks are common across the three protocols (Figure 2C). We observed that even the NPOP-seq can capture a significant fraction of genes targeted by proteins, while missing ~19% of the total identified genes by the other two versions of POP-seq (Figure 2C).

**Figure 2.**
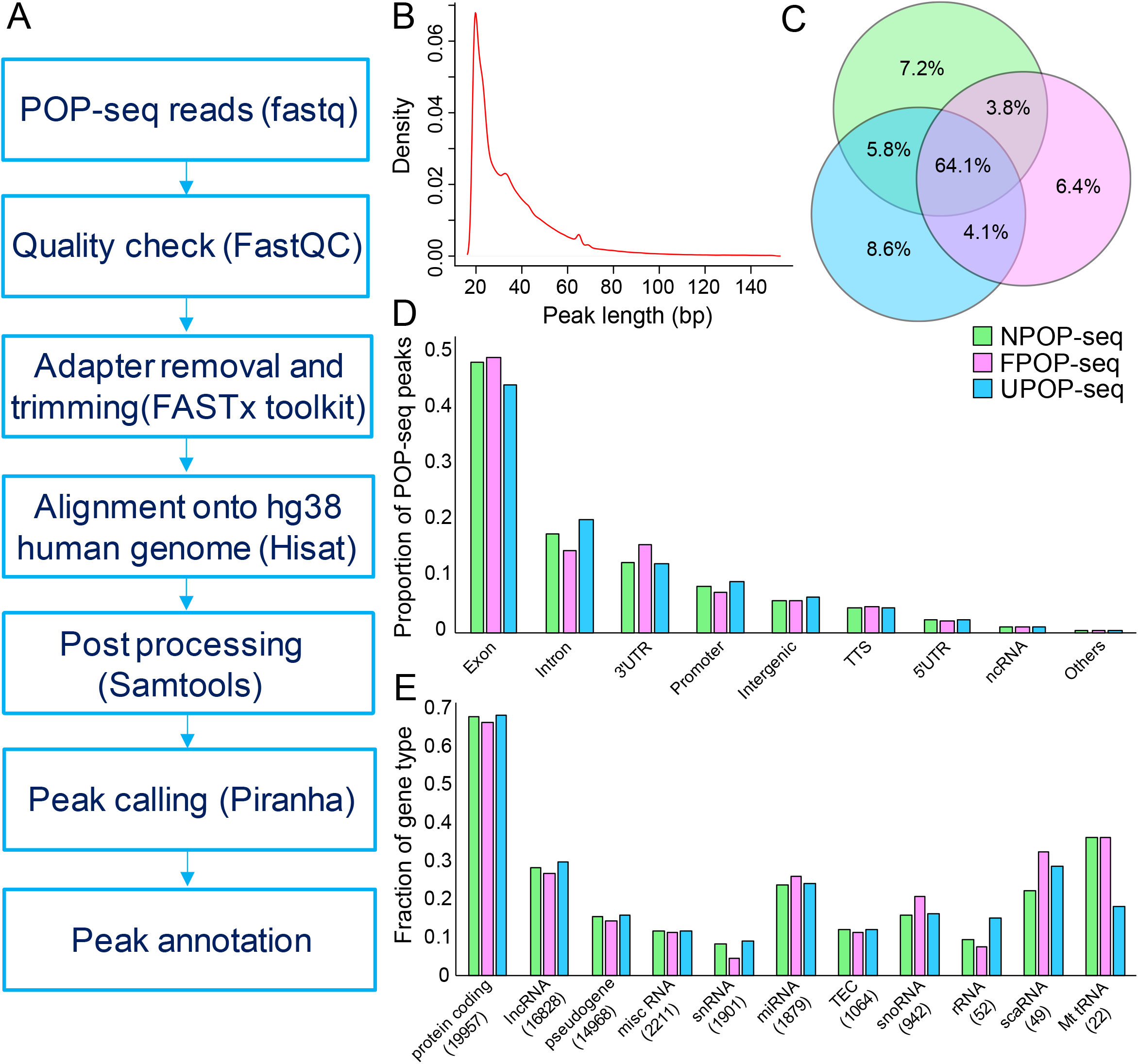
Statistical analysis of POP-seq dataset. (A) Workflow for POP-seq data processing and downstream analysis (B) A density plot showing the distribution of POP-seq peaks length (bp) (C) Venn diagram showing the overlap of genes exhibiting POP-seq peaks across the protocols. (D) Proportion of POP-seq peaks in genomic elements (E) Fraction of gene types captured by POP-seq.

Next, we examined the distribution of POP-seq peaks across transcript regions annotated by HOMER and observed that majority of the peaks were mapped to the exons (~48%) while a relatively lower proportion were mapped to intronic regions (~19%) and 3’UTR’s (~16%) (Figure 2D). Our observation supporting the higher proportions of the peaks in exonic regions was in agreement with the previous reports in MCF7 and HEK293 cell lines^18,19^. Next, we examined the fraction of gene types exhibiting the POP-seq peaks which revealed that majority of the genes (~67.3%) mapped to protein coding followed by ~28.4 % lncRNA and ~ 18.6 % snoRNA with respect to the total genes annotated in the human reference genome (hg38) (Figure 2E). These proportions were generally consistent across all the POP-seq protocols.

### POP-seq illustrates a significant enrichment for protein-RNA interactions

To rigorously evaluate whether POP-seq identifies any non-RBP binding events, we performed three comprehensive analyses as summarized below (i) To estimate non-RBP interactions, we compared the POP-seq peaks with ChIP-seq data of 67 proteins and CLIP-seq data of 79 proteins available for K562 cells (from ENCODE project). We observed a significantly higher overlap (p value < 2.2e-16) of POP-seq peaks with CLIP-seq peaks compared to ChIP-seq peaks as shown in Figure S2A, (ii) To estimate the protein-DNA interactions (false positives) that could be captured by POP-seq, we systematically compared the POP-seq peaks (and 5 random peak profiles separately) with the binding profile of 18 proteins for which both ChIP-seq and CLIP-seq data was available in K562 cells (from ENCODE project). Our results showed a significant enrichment (Odds ratio > 20 averaged across 5 random controls, p value < 2.2e-26, Fishers Exact test) of POP-seq signals overlapping with the CLIP-seq profile than the ChIP-seq profile of these 18 proteins (Figure S2B), indicating that POP-seq peaks are enriched for protein-RNA interactions. Among the 18 tested proteins, NONO (Non-POU domain-containing octamer-binding protein), which is known to bind both DNA and RNA^30,31^, expectedly demonstrated relatively similar significance of binding to both DNA and RNA compared to random locations. Overall, our analysis shows that irrespective of POP-seq protocols, signals are underrepresented in ChIP-seq data while overrepresented in the CLIP-seq data indicating a clear enrichment for RNA binding events compared to publicly available protein-DNA maps. (iii) To estimate the ribosomal protein interactions captured by POP-seq, we compared the POP-seq peaks with publicly available ribo-seq data in K562 cells. Our analysis showed that ~20% of the total POP-seq peaks (with peak length ≤50 bp) exhibited 50% end-to-end overlap with the ribo-seq peaks indicating that some ribosomal protein-RNA interactions are captured by POP-seq.

### Comparison of POP-seq data with Formaldehyde and UV crosslinked RBPs reveals high quality of POP-seq peaks

POP-seq is a technique which provides occupancy levels for proteins on a global transcriptome-wide scale. The functionality of thousands of binding sites that are generated resulting from this method could correspond to scores of RBPs and RNP complexes. Since there are no large-scale assays currently available to validate the RNA-protein interaction sites globally, and the low throughput assays will not cover large number of interaction sites, therefore, we employed orthogonal methods using publicly available data for 24 formaldehyde crosslinked RBPs^32^ with respect to five random non-peak files (See methods) and eCLIP profile of 97 RBPs^28^ in the K562 cells from ENCODE (see Methods). Our analysis indicates a significant enrichment of POP-seq peaks in both CLIP-seq (Figure 3A for top 24 RBPs, supplementary Figure S3) and fRIP-seq profile (Figure 3B) of individual RBP’s compared to 5 random non-peak profiles. Confusion matrix for this analysis is documented in supplementary Table S1 and S2. Overall, our analysis shows that POP-seq can recover high quality peaks corresponding to specific RBPs identified from individual crosslinking protocols.

**Figure 3.**
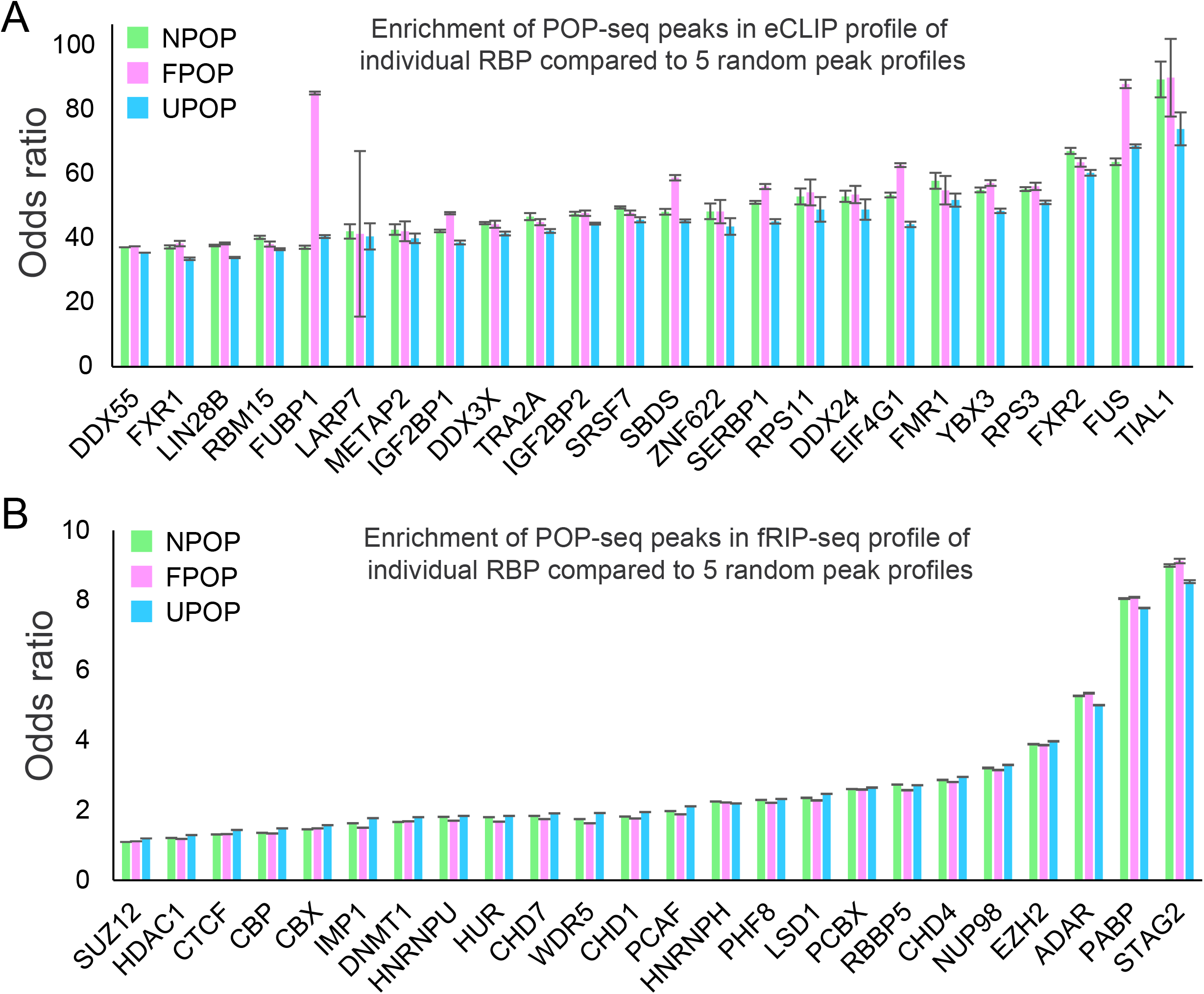
Comparison of POP-seq peaks with RBP centric peaks derived from orthogonal assay. Bar plot showing the enrichment of POP-seq peaks in (A) eCLIP and (B) fRIP-seq profiles of individual RBP in K562 cells, compared to 5 random peak profiles and statistically tested using Fisher’s exact test.

### CRISPR knock out RNA-sequencing data of RBPs supports the functionality of POP-seq peaks

Development of targeted genome editing using CRISPR has revolutionized the genomic research^33^, particularly to understand the molecular mechanism involved in gene regulation and expression^34,35^. A recent study by our research group demonstrated that the functional relevance of protein-RNA interactions can be estimated by the expression of the exons upon perturbation of RBP binding site in their neighborhood using CRISPR-Cas9 system^36^. Therefore, we aimed to interrogate the functional impact of POP-seq captured protein-RNA interactions by its systematic comparison with the eCLIP profile of RBP’s, for which knockout data is publicly available in ENCODE project^28,37^ For this analysis, we used two RBPs; a) DGCR8 (DiGeorge Syndrome Critical Region 8), which is involved in microRNA processing and is implicated in the pathogenesis of cancer^38,39^ and b) IGF2BP1 (insulin like growth factor 2 mRNA binding protein 1) which is a critical post-transcriptional regulator of various mRNA involved in cancer progression^40^. First, we identified the POP-seq peaks from the individual protocol that showed >50% base-to-base overlap with eCLIP profile of respective RBP (obtained from ENCODE^28^). Next, we extracted the expression levels of exons ‘proximal’ (<1000 bp) to overlapped peaks from CRISPR knock out data set (Material and Methods). We observed that the cumulative expression level of ‘proximal’ exons was significantly dysregulated with respect to the nontargeting control. We observed that there was a significant reduction in the expression of ‘proximal’ exons in DGCR8 KO and a significant increase in IGF2BP1 KO with respect to their non-targeting CRISPR control (Figure 4). However, there could be alternative hypotheses such as the contribution of other binding sites from same or different RBPs that could account for the compensatory effects in expression levels in our CRISPR analysis, which could explain why not all the proximal exon levels are altered. More importantly, RBP binding does not always alter the expression of the target exon/transcript but instead may contribute to editing, structure and localization of bound RNA. However, despite these alternate possibilities, it is promising to observe that the loss of binding sites has a significant impact on the target exons. Overall, this analysis suggests that POP-seq can capture the functionally relevant protein bound sites and indicates that the dysregulation of exons proximal to the functional binding site occur in an RBP dependent manner.

**Figure 4.**
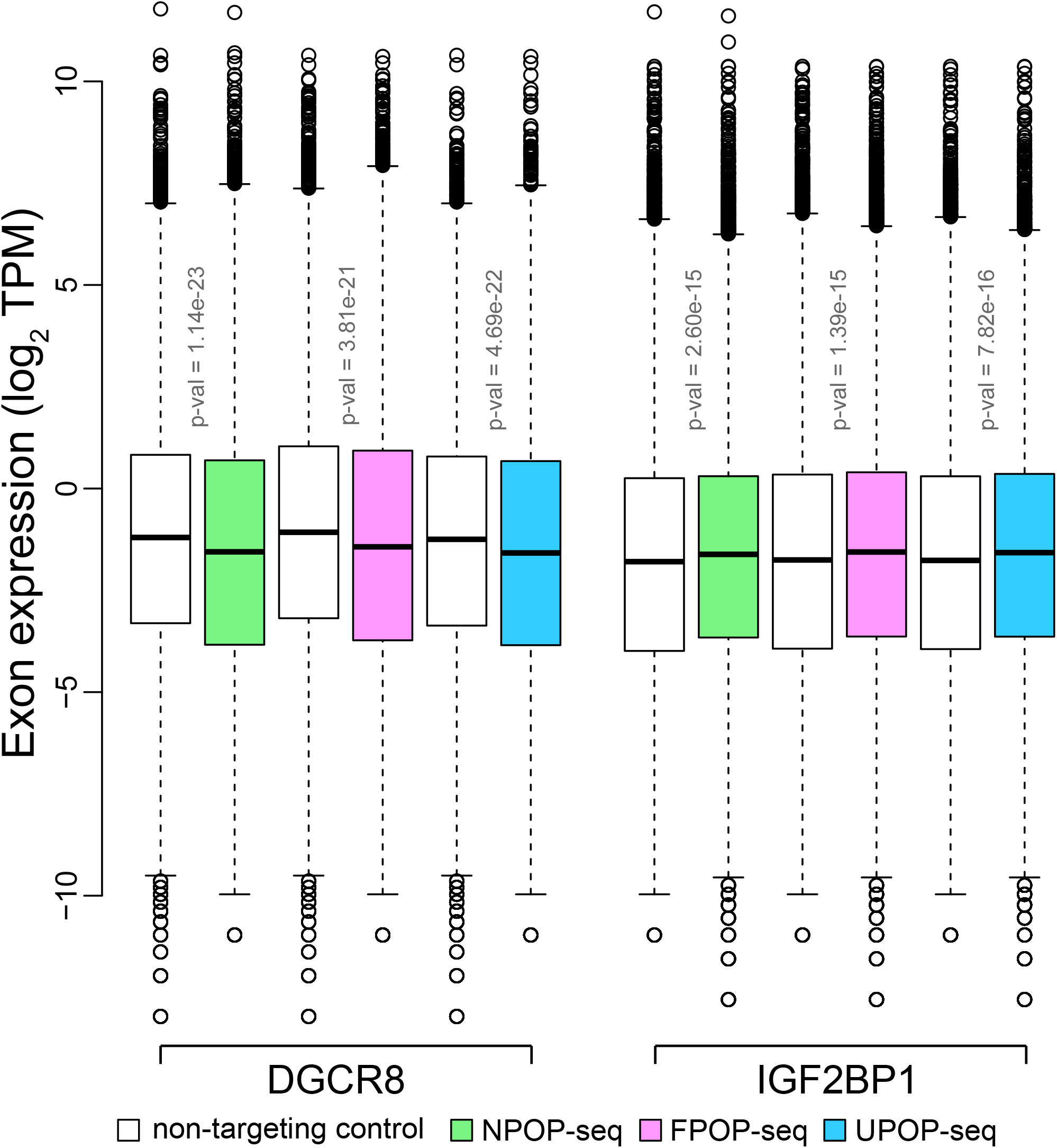
Comparison of POP-seq data with CRISPR knock out RNA-sequencing data of RBPs. Box plot showing the cumulative exon expression levels proximal (<1000 bp) to POP-seq peaks overlapped (50% end to end peak overlap) with the eCLIP profile of DGCR8 and IGF2BP1 in K562 cells (ENCODE project). The exon expression levels of respective RBP CRISPR Cas9 knock out was compared with respective non-targeting Crispr control and statistically tested using Wilcoxon test.

### POP-seq supports the protein-RNA binding sites across regulatory gene families

To obtain a detailed perspective of the peaks captured by POP-seq, we examined the occurrence of the protein-RNA binding sites across different regulatory genes, classified as RNA binding protein, ENO1 (Enolase1)^41^, lncRNA MALAT1 (metastasis associated lung adenocarcinoma transcript 1)^42^ and a transcription factor, Jun^43^. ENO 1 is a crucial glycolytic enzyme involved in cell growth and is also reported as an oncogene that promotes metastasis by facilitating cell proliferation in multiple cancers including colorectal, lung, and prostate cancer^44–48^. We observed the abundance of protein bound pockets in the genomic loci of ENO 1 across the POP-seq protocols (Figure 5A). We also investigated the protein-RNA interactions in MALAT 1, a highly conserved lncRNA that governs a variety of functions including regulation of gene expression, alternative splicing, neural development and vascular growth^49–51^. Several studies report the abundant expression of MALAT1 in multiple cancers such as lung cancer, bladder cancer, breast cancer, colorectal cancer and others^49,52–55^. Therefore, identification of regulatory sites targeted by proteins in MALAT1 is crucial for understanding its pathogenesis in cancers. Our data suggest that all the three protocols can capture the abundance of regulatory sites targeted by proteins in the genomic boundary of MALAT1 (Figure 5B). We observed relatively enhanced signals for protein occupancy sites in MALAT1 locus in UPOP-seq compared to the NPOP-seq and FPOP-seq (Figure 5B). To our observation, NPOP-seq also captured the physiologically relevant protein-RNA interactions in MALAT1 suggesting its application for unbiased protein occupancy profiling (Figure 5B).

**Figure 5.**
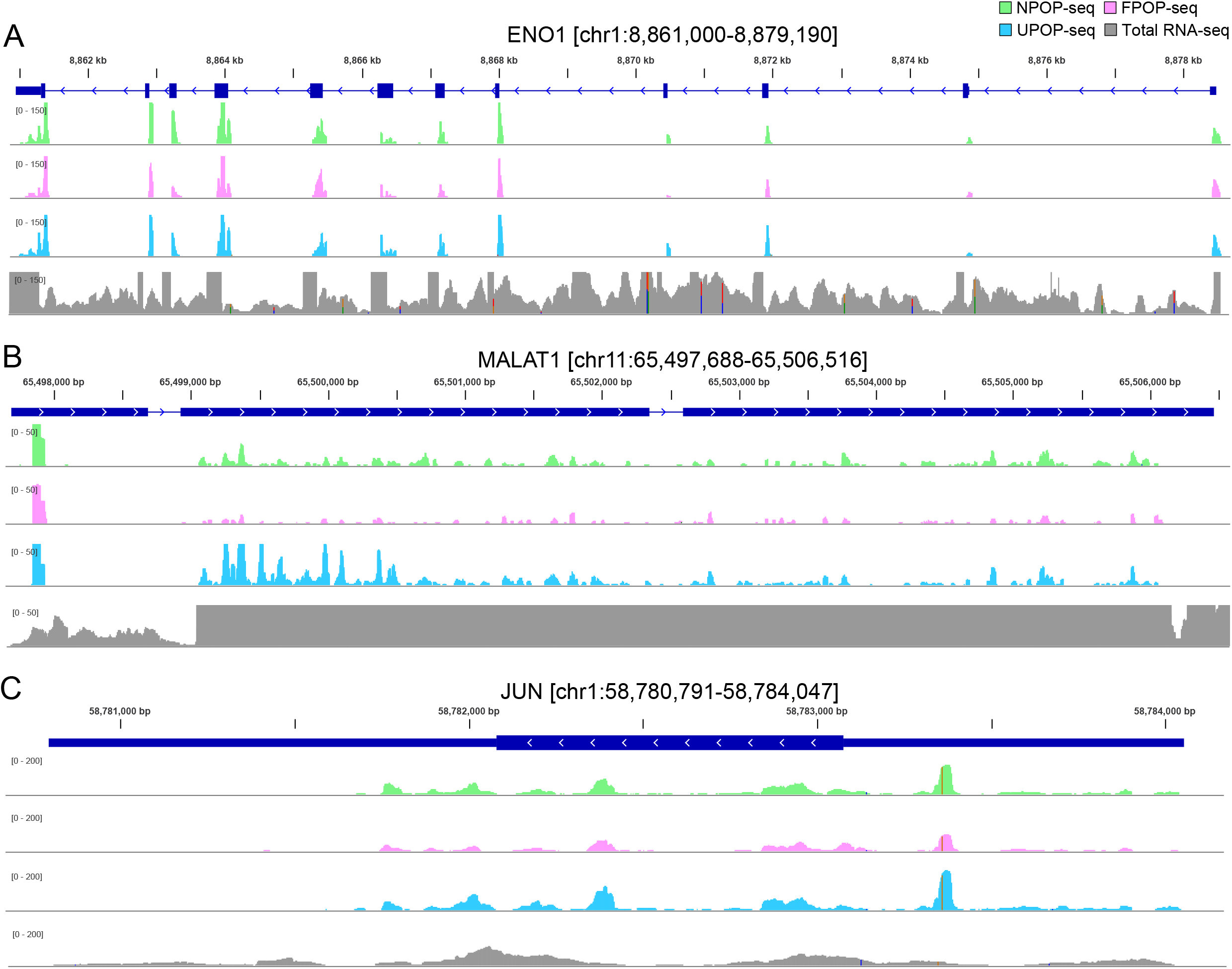
Genomic tracks showing the POP-seq peaks spanning the genomic loci of regulatory genes; (A) RBP – Enolase1 (ENO1) (B) LncRNA – MALAT1 and (C) Transcription factor – Jun. Total RNA-seq data (from K562 cells) is included in the track as a background control (Y-axis adjusted to POP-seq scale).

We further explored the regulatory binding sites in AP1 transcription factor subunit ‘Jun’ which is a proto-oncogene actively involved in cell proliferation, apoptosis, inflammation and carcinogenesis^56–60^. We found the occurrence of POP-seq peaks in Jun’s genomic boundary across all the three protocols (Figure 5C) consistent with the observations in ENO1 and MALAT1. Overall, the results demonstrate that POP-seq can capture the protein-RNA interaction sites across the regulatory gene families.

### POP seq identifies germline and somatic variants that potentially contribute to altered post transcriptional regulation

Single nucleotide polymorphisms (SNPs) are reported as the most common form of somatic variations and are widely associated with metabolism, cell cycle regulation and DNA mismatch repair^61,62^. In past years, SNPs has emerged as a potential diagnostic biomarker for several cancer types^63–66^. Therefore, it is imperative to investigate the somatic variations arising due to SNPs and their effect on transcriptome wide protein RNA interaction sites.

In order to detect the somatic variations captured by POP-seq, we calculated the proportion of known somatic variations (see Materials and Methods) occurring in the equivalent genomic loci of the peaks. We tested the enrichment of genomic variations from GWAS catalog^67^ and Ensembl Variation database^68^ (PhenVar, ClinVar and somatic variations) in POP-seq peaks than expected by chance (i.e. 5 random non-peak profiles) using Fisher’s exact test. For this analysis, random ‘non-peak’ files were generated as described previously. Our results indicate a significant enrichment for each SNP cohort with relatively lesser enrichment for GWAS SNP (averaged odds ratio=1.45, 1.42, 1.35 for NPOP, FPOP and UPOP-seq respectively, p value<2.2e-16). Similar test for other genomic variations including PhenVar, SomaticVar and ClinVar indicated relatively higher enrichment (Odds ratio ~ 22 averaged across 5 random controls for each cohort, p value < 2.2e-16, Fishers Exact test) in POP-seq peaks compared to non-peaks (Figure 6A). This observation provides support for the enrichment of both germline and somatic SNPs including those reported with clinical significance to be prominent on protein RNA interaction sites, implying the need for deeper understanding of their functional consequences. Indeed, we identified the occurrence of two clinically relevant genetic risk loci from GWAS; rs45461499 in CDC20 (Cell-division cycle protein 20) and rs7578199 in HDLBP (High Density Lipoprotein Binding Protein) genes that are reported in acute and chronic lymphoblastic leukemia respectively^69,70^ (Figure 6B and C) to harbor protein-RNA interaction sites. Thus, our implementation of POP-seq in K562 cells demonstrate a novel and robust approach to elucidate the occurrence of somatic variants in leukemic patients.

**Figure 6.**
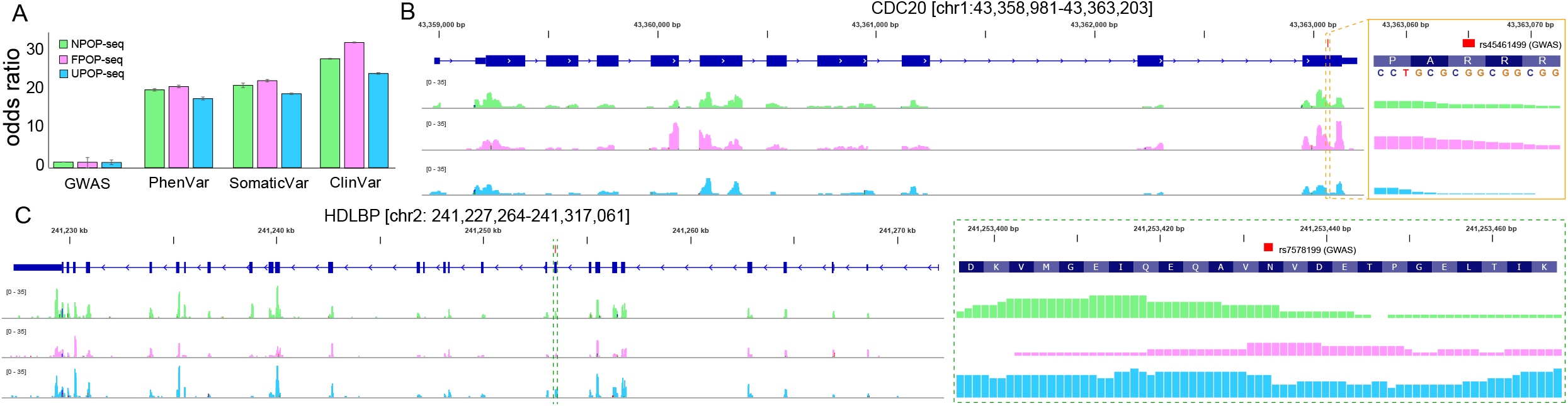
Somatic variation captured in POP-seq peaks. (A) Bar plot showing the enrichment (odds ratio) of genomic variations (GWAS, PhenVar, ClinVar and somatic variations) in POP-seq peaks, statistically tested using Fisher’s exact test. Genomic tracks showing the POP-seq peaks spanning the genetic risk loci (GWAS SNP) associated to (B) acute myeloid leukemia in CDC20 and (C) chronic myeloid leukemia in HDLBP.

### Highly expressed lncRNAs exhibit abundance of POP-seq peaks in K562 cells

Long non-coding RNAs (lncRNAs) have been widely documented with diverse roles in the transcriptional and post-transcriptional regulation of gene expression^71,72^. Aberrant expression of lncRNA is associated with the pathogenesis of various diseases including cancer^73^ and have been profoundly recognized as pivotal targets in cancer therapeutics^74^. However, the mechanism underlying lncRNA regulation is not well elucidated^75^. Therefore, we speculated that associating the expression of lncRNA with the occurrence of POP-seq peaks would provide an insight into the transcriptome wide regulation of lncRNAs. We observed that the highly expressed lncRNAs exhibit abundance of regulatory binding sites in K562 cells (Figure 7A, Materials and Methods), suggesting that lncRNAs dynamically interact with RBPs in pathological conditions. In general, highly expressed RNAs are expected to be more available for binding by proteins and therefore exhibit higher RNA binding events. Therefore, we tested the association between the expression levels (high and low) with the number of POP-seq peaks per unit length for both lncRNA and non-lncRNA genes across the technical replicates. We found that the replicates exhibited a reproducibility in the trend (Figure S4A), irrespective of the POP-seq protocols. To test whether such a trend can also be observed for non-lncRNA, we carried out the same analysis across the replicates. We observed that there is tendency for even non-lncRNAs to exhibit higher expression with more binding sites, however the trend is not as robust with lower significance (Figure S4B) compared to lncRNA as shown in Figure S4A.

**Figure 7.**
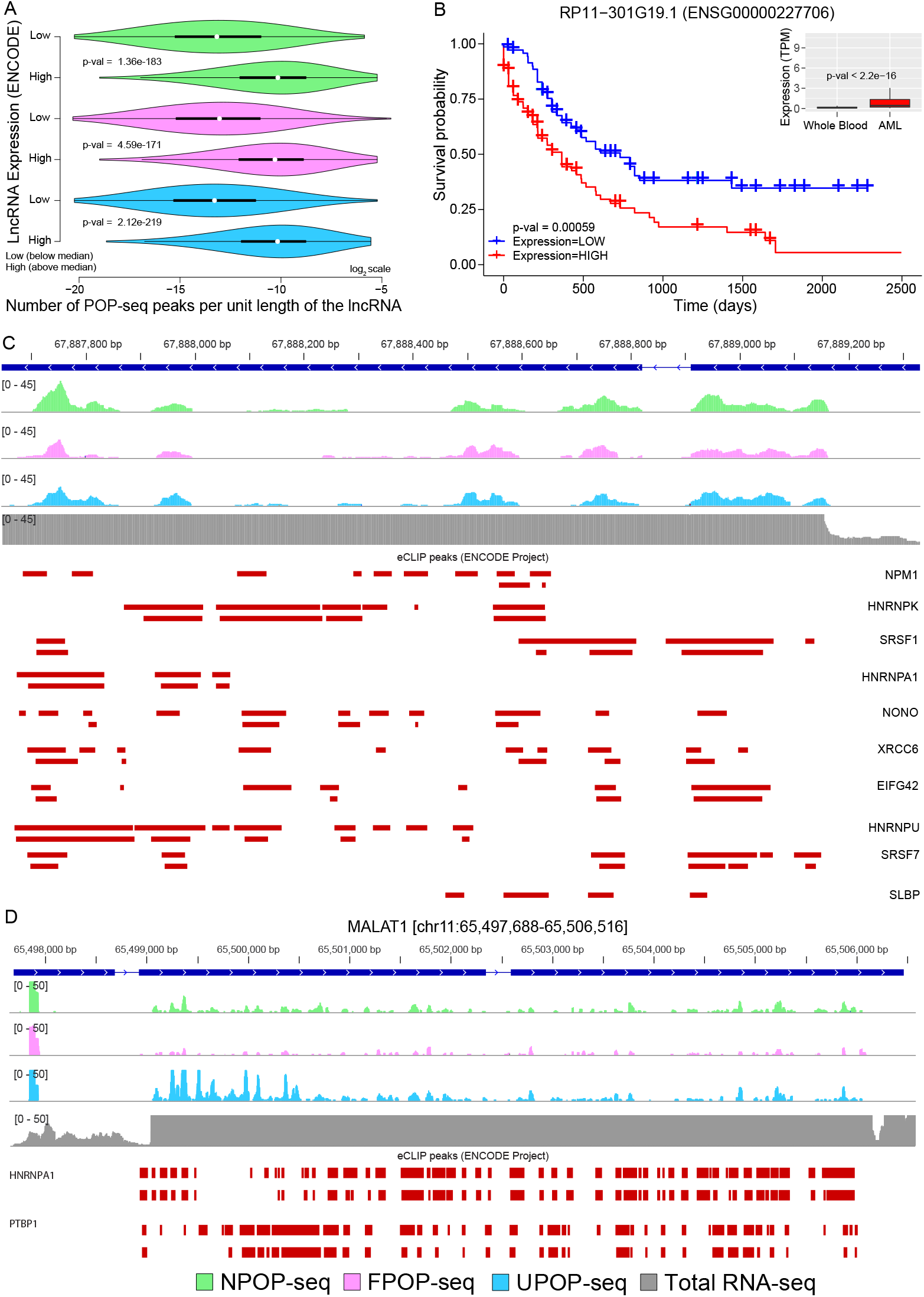
Comparative analysis of POP-seq peaks with lncRNA expression. (A) Violin plot showing the number of POP-seq peaks (normalized per unit length of the lncRNA) binned in low and high expression group of LncRNAs in K562 cells. The difference in normalized peak counts between the two groups was statistically tested using Wilcoxon test. (B) Kaplan-Meier plot showing the association of lncRNA RP11-301G19.1 expression with the survival of AML patients (in days). An inset showing the expression of RP11-301G19.1 in whole blood (GTEx) cohort and AML patients (from TCGA) where the difference between the two groups was statistically tested using Wilcoxon test (C) Genomic tracks showing the distribution of POP-seq variant peaks in RP11-301G19.1 along with the eCLIP profile of highly expressed RBPs in K562 cells. (D) Genomic tracks showing the distribution of POP-seq peaks in MALAT1 along with the eCLIP profile of HNRNPA1 and PTBP1 in K562 cells. Total RNA-seq data (from K562 cells) is included in the track as a background control (Y-axis adjusted to same POP-seq coverage scale).

Several studies have reported that the majority of the lncRNAs exhibit a ‘sponge effect’ to titrate the abundance of regulatory proteins such as RBPs in a cell type specific manner^76–78^. These studies proposed that lncRNA sponges can extensively rewire the post-transcriptional gene regulatory networks by altering the protein–RNA interaction landscape in cell-type and phenotype specific manner. Based on this finding we tested the hypothesis that highly expressed lncRNAs could sponge the regulatory RBPs that would further provide an insight into lncRNA mediated regulation of RBPs in disease context. We uncovered RP11-301G19.1, a highly expressed lncRNA in leukemia^79,80^, to illustrate the ‘sponge effect’ in K562 cells. We found an abundance of this lncRNA in AML patients from TCGA^81^ compared to its expression level in normal whole blood from GTEx cohort^82^ (Figure 7B, inset). In order to predict whether the expression of this lncRNA contributes to the survival of AML patients, we employed Kaplan-Meier survival analysis^83^ for RP11-301G19.1. We found that this lncRNA exhibits a significant (False Discover Rate (FDR) < 0.00059) prognostic impact in AML patients (Figure 7B). Next, we interrogated the regulatory sites captured by POP-seq in the genomic loci of RP11-301G19.1 and observed a consistent occurrence of peaks across all POP-seq protocols. Our results demonstrate that majority of the POP-seq peaks overlapped with the eCLIP profile of multiple RBPs (Figure S5) which further supports the “sponge effect” for RP11-301G19.1. A subset of highly expressed RBPs in K562 cells illustrates a general agreement of their eCLIP profile with POP-seq peaks in RP11-301G19.1 gene (Figure 7C). To further elaborate the sponge effect, we investigated a well-studied lncRNA MALAT1 which has been shown to interact with numerous RBPs^84–88^. As illustrated in Figure 7D, we observed that POP-seq captured peaks were sparsely distributed along the length of MALAT1 in contrast to the fairly uniform distribution of total RNA-seq reads showing higher expression of MALAT1 in K562 cells. We also included a track of ENCODE eCLIP peaks for RNA binding proteins HNRNPA1 and PTBP1 with known binding to MALAT1^84^ shown in Figure 7D. The results suggest that HNRNPA1 and PTBP1 are most likely being sponged by highly expressed lncRNA MALAT1 in K562 cells. Additionally, we also observed that other RBPs with transcriptome wide interaction maps available from ENCODE project exhibited several binding sites overlapping with POP-seq peaks along the length of the MALAT1 and NEAT1 lncRNAs (Supplementary Figure S6A and B). Overall, the results suggest that POP-seq can capture the occupancy sites of RBPs on lncRNAs and thus advance our understanding of lncRNA regulation in diseases.

## Discussion

Protein-RNA interaction is a vital phenomenon regulating crucial transcriptional and post-transcriptional processes starting from intercalation of the DNA-RNA juncture to RNA metabolism, translation and decay^89,90^. RNA binding proteins have been widely recognized as the key regulatory proteins for these processes^91^. Although the past few decades have seen a surge in the number of methods for capturing protein-RNA interaction sites occupied by RBPs on a transcriptome-wide scale^18,19,21^. Majority of these protocols employ UV or formaldehyde cross linking and poly-A pull followed by sequencing of RNA. However, this excludes their application to non-polyadenylated RNAs and so far, a systematic analysis to depict the effect of crosslinking on the captured interaction sites has not been investigated.

In this study, we present POP-seq to assess the protein bound RNA fragments using trizol based phase separation. POP-seq can uniquely map the protein bound RNA pockets in a transcriptome wide manner. We implemented three versions of POP-seq to generate an unbiased profile of protein-RNA interactions in K562 cells. We demonstrated that POP-seq captured peaks generally agreed with the eCLIP profile of several RBPs. The abundance of protein-RNA interactions captured by POP-seq was mostly observed in exonic followed by intronic regions, with consistent overlap across the POP-seq protocols in K562 cells. Further analysis of publicly available CRISPR KO dataset of RBPs from ENCODE project^92^, illustrate that POP-seq can capture the functionally relevant protein bound pockets implying that the dysregulation of exons proximal to the functional binding site occur in RBP dependent manner. POP-seq also enables the identification of clinically relevant somatic variations associated to leukemia. Further, POP-seq provides a comprehensive evidence of the potential protein binding sites in most of the regulatory gene families such as TFs, RBPs and lncRNAs.

Since ribosomal protein complexes constitute the basic translation machinery and hence are expected to be highly abundant in cells. To evaluate the prevalence of ribosomal protein occupied sites in our data, we compared POP-seq peaks with publicly available ribo-seq data^93^ in K562 cells. We observed that ~20% of the total POP-seq peaks (with peak length ≤50 bp) exhibited 50% end-to-end overlap with the ribo-seq peaks and this fraction was significantly reduced with increase in % peak overlap as shown in supplementary Table S3. These observations suggest that a fraction of POP-seq peaks correspond to the regions occupied by ribosomal complexes indicating a potential for further optimization of the protocol to enhance the stringency for selectively capturing sites occupied by regulatory RBPs under physiological conditions

Interestingly, POP-seq provides evidence for the ‘sponge effect’ depicted by lncRNA on multiple RBPs thus advancing our understanding of the post-transcriptional regulatory mechanism controlled by lncRNAs interacting with RBPs in cancer. In summary, POP-seq is a cost-effective and robust approach to elucidate the binding sites of proteins in a transcriptome-wide manner. Thus, it should stand as a generic framework for mapping the global protein-RNA interactions, widening the scope and application of this technique to primary tissues for rapid profiling of protein occupancies.

## Materials and methods

### Cell culture

K562 cells were obtained from the American Type Culture Collection (ATCC). Cells were cultured in Dulbecco’s minimal essential medium (DMEM, Gibco) supplemented with 10% heat-inactivated fetal bovine serum (FBS, Atlanta Biologicals) along with 1% antibiotics (penicillin 5000 Units/ml, Streptomycin 5000 μg/ml). All cells were maintained at 37°C and 5% CO_2_ in a humidified incubator and fresh media was replenished every alternate day until confluent.

### Crosslinking

Cells were cultured in T-175 flask until a maximum of 90% confluency was reached. A total of 20 million cells per replicate of each sample were used for UV, formaldehyde, and no-crosslinking approaches. Cells were washed twice with 1X PBS, the supernatant was removed by pipetting and cells were resuspended in 1X PBS. For UV crosslinking, cells in PBS suspension were transferred to 100 mm dishes and UV irradiated at 254 nm with 400 mJ/cm^2^ dosage (UV Stratalinker 1800). Immediately after crosslinking, cells were collected in 15 ml tubes and pelleted at 1500 rpm for 5 min. Supernatant was discarded and cells were lysed in 1 ml trizol reagent (Life Technologies) by pipetting up and down to obtain a homogenous cell lysate.

For Formaldehyde crosslinking, cells were first washed in 1X PBS twice until all the media is removed. Next, cells were resuspended in 30 ml PBS and crosslinked with 0.5% formaldehyde for 10 min by gentle shaking at room temperature. To stop crosslinking, 1 M Glycine was added to the cell suspension for 5 min by gentle shaking at room temperature. Cells were pelleted down at 1500 rpm for 5 min, lysed in 1 ml trizol and homogenized by pipetting up and down. For noncrosslinked samples, cells were pelleted down, washed in 1X PBS and immediately lysed in trizol for phase separation.

### Guanidinium Thiocyanate–Phenol–Chloroform (TRIZOL) extraction

Trizol lysed cells were incubated at room temperature for 5 min to dissociate the weak RNA-protein interactions. Phase separation was achieved by adding 200 μl chloroform and thoroughly mixed by vortexing, followed by incubation at room temperature for 5 min. Samples were then centrifuged at 12000 g for 10 min at 4°C to obtain three phases: aqueous phase (top), interphase (middle) and organic phase (bottom). The aqueous layer was discarded by pipetting and the organic layer was discarded by passing the tip through the interphase leaving behind up to 100 μl of the organic layer. The interphase was resolubilized in trizol followed by phase separation with chloroform three times. After the third phase separation, the interphase was precipitated by adding 1 ml methanol, spun down to remove supernatant containing methanol.

### POP-seq strategy

Following the trizol lysis and phase separation, POP-seq was implemented on the three versions in replicates. Interphase pellet was subjected to RNase A/T1 (Thermo Scientific) degradation in RNase buffer (10 mM Tris-HCl, pH 7.5, 300 mM NaCl and 5 mM EDTA, pH 7.5). 2 μg of RNase A/T1 mix was added to the interphase pellet, mixed by pipetting, and incubated at 37°C for 1 h. Interphase-RNase mixture was resolubilized in trizol reagent to recover the RNA-protein complexes. The aqueous and organic layers were discarded as described previously. Interphase pellet was precipitated in 1 ml methanol. Next, the interphase was mixed with Proteinase K (Ambion) in appropriate buffer (0.1M NaCl, 10 mM Tris-HCl, pH 8, 1 mM EDTA, 0.5% SDS and 200 μg/ml proteinase K) and incubated at 50 °C for 2 h. After proteinase K digestion, free RNA was recovered from the aqueous layer by trizol extraction as described previously.

Purified RNA concentration was estimated using Nanodrop and up to 1 μg of RNA was incubated with DNase I, 1U (Thermo Scientific) at 37°C for 30 min to remove any traces of DNA contamination. 1 μl of 50 mM EDTA was added to the reaction mixture and incubated at 65°C for 10 min to terminate the reaction. RNA was purified using trizol extraction from the aqueous layer. At this point, r-RNA depletion was performed with 1 μg input RNA using Ribocop kit (Lexogen) as per manufacturer instructions. Further, the ends of r-RNA depleted RNA were modified by treating with Calf intestine alkaline phosphatase (CIAP, Invitrogen) and T4 polynucleotide kinase (T4 PNK, Thermo Scientific) as per manufacturer protocol. The end modification enabled the library preparation of these RNA fragments.

### RNA integrity, library preparation and sequencing

RNA purity and concentration were assessed at each step using Nanodrop, based on the absorbance ratio 260/280 >2. RNA integrity was evaluated using Agilent 2100 bioanalyzer system. At least 50 ng of r-RNA depleted RNA was used to generate sequencing libraries using the True-seq small RNA library prep kit (Illumina). All libraries were barcoded and sequenced in parallel on a Next-seq platform for 400 million reads to obtain 75 bp single end reads.

### Data processing and statistical analysis of POP-seq peaks

We implemented NGS data processing pipeline to facilitate the analysis of the POP-seq data as shown in Figure 2A. Firstly, we investigated the quality of sequenced reads using FASTQC (http://www.bioinformatics.babraham.ac.uk/projects/fastqc/) and deployed FASTX-toolkit (http://hannonlab.cshl.edu/fastx_toolkit/) for removal of adapters and low quality read fragments wherever applicable. Next, we aligned the high quality reads onto human reference genome (GRCh38.p12) using HISAT^94^ followed by post processing using Samtools^95^. To ensure the reproducibility between the replicates, we compared the aligned reads per 10 kb genome using deepTools ‘plot correlation’ module^26^. We employed Piranha^96^ for peak calling with default parameters and obtained the resulting POP-seq peaks in bed format. Source code of the POP-seq data processing pipeline is accessible at GitHub (https://github.com/Janga-Lab/POP-seq). We merged the replicate bed files of respective POP-seq protocols and used several tools such as bedtools^97^, HOMER^29^ (annotatePeaks.pl), and R (https://www.r-prqiect.org/) for annotation, statistical testing and other downstream analysis. We also downloaded the publicly available formaldehyde RNA ImmunoPrecipitation (fRIP-seq) data^32^ for 24 RBPs from GSE67963 and raw ribo-seq data^93^ from GSE125218, both generated in K562 cells and processed them using the same pipeline and identified the peaks. Next, we computed the fraction of POP-seq peaks overlapping with the identified peaks from both the datasets independently using bedtools^97^.

Similarly, we downloaded the eCLIP^98^ profiles of 97 RBPs from ENCODE project^28^ and concatenated the unique coordinates into a bed file. Resulting coordinates of the binding sites of RBPs were compared with the POP-seq peaks using bedtools^97^. All POP-seq data have been deposited under Gene Expression Omnibus (GEO) accession number GSE142460. We also downloaded the raw FATSQ reads of total RNA (accession no. ENCLB822JYE from ENCODE project) and processed using the standard NGS pipeline as described in below section.

### Comparison of POP-seq peaks with publicly available CLIP-seq and ChIP-seq data

We downloaded the ChIP-seq and CLIP-seq profile of available proteins for K562 cells from ENCODE project^28^. We computed the overlap of POP-seq peaks with CLIP-seq peaks and ChIP-seq peaks using bedtools^97^. This dataset also includes 18 proteins for which both ChIP and CLIP-seq data is available for K562 cells. We generated 5 random peak profiles for each POP-seq protocols using bedtools ‘shuffle’ function. Each random peak profile contains equal but “non-peak” locations (with consistent peak width distribution), within the gene boundary. Then, we compared the POP-seq derived peaks (and the random peaks separately) with the ChIP and CLIP-seq profile of 18 proteins. We employed Fisher’s exact test to estimate the level of significance for the enrichment of POP-seq in protein bound DNA(ChIP-seq)/RNA(CLIP-seq) locations compared to non-peaks.

### Integrated data analysis of POP-seq peaks with the CRISPR knock out studies

CRISPR/Cas9 Knock Down (KD) followed by expression profiling on several RBPs have been conducted as part of the ENCODE project^28^ to facilitate the understanding of downstream biological processes associated to loss of function of the respective RBP. We downloaded the raw RNA-sequencing dataset of CRISPR experiments of DGCR8 (n=6), IGF2BP1 (n=2) where gRNAs were used to deplete the functional form of RBPs and their non-specific CRISPR control (n=8) in K562 cells^92^. We processed the raw sequencing reads using standard NGS data analysis pipeline (as described previously). Briefly, we filtered the low-quality reads (Phred Score < 30) using FASTQC tool and aligned them onto human reference genome (hg38) using HISAT^94^. After post processing (using samtools), we used StringTie^99^ to quantify the expression levels in Transcripts Per Million (TPM) reads for all the genes annotated in human genome (hg38). Thereafter, we used an ad-hoc script to calculate the exon levels from the resulting gtf files of StringTie and converted the resulting files into an exon expression matrix.

To further investigate the functional relevance of protein-RNA interactions, we identified the POP-seq peaks from the individual protocols that exhibited at least 50% overlap with an eCLIP profile of the respective RBP (as described previously). Next, we extracted the expression levels of exons, proximal (<1000 bp) to overlapped peaks, from exon expression matrix of the knock down experiments. We compared the distribution of the expression levels of these proximal exons in non-targeting control to that in respective KD and statistically examined the condition specific expression differences using Wilcoxon test^100^.

### Identification of somatic variants in protein-RNA interacting sites

Several studies suggest that single nucleotide variations (SNVs) play an important role in gene regulation via riboSNitches’^101^ i.e. by altering RNA secondary structure or TAM (Transcript associated mutation) that further contribute to transcriptome complexity in higher eukaryotes. Therefore, it is imperative to investigate the genomic variations occurring in protein-RNA interaction sites identified by POP-seq protocols. Hence, we downloaded the somatic variants reported in the GWAS catalog^67^ and Ensembl Variation database^68^ including phenotype and clinically associated somatic variations (ftp://ftp.ensembl.org/pub/release-97/variation/vcf/homo_sapiens/). In order to detect the somatic variations captured by POP-seq, we tested the enrichment of genomic variations from GWAS catalog^67^ and Ensembl Variation database^68^ (PhenVar, ClinVar and somatic variations) in POP-seq peaks than expected by chance (i.e 5 random non-peak profiles) using Fisher’s exact test. For this analysis, random ‘nonpeak’ files were generated as described previously. To gain disease specific understanding of the role of SNPs in impacting protein-RNA interactions, we also investigated the POP-seq peaks overlapping with SNPs associated with leukemia (from GWAS) and generated genomic tracks in Integrative Genomics Viewer (IGV)^103^ for CDC20 and HDLBP genes.

### Comparative analysis of POP-seq peaks across lncRNAs and their association with lncRNA expression

We mapped the protein-RNA interaction sites captured by POP-seq protocols onto known lncRNAs using bedtools. For each lncRNA, the number of POP-seq peaks normalized by respective gene length was calculated. To obtain the expression levels, we processed the raw RNA sequencing dataset (paired end reads, n=5, in replicates) of K562 cells from ENCODE using a standard NGS data analysis pipeline described earlier and generated a gene expression matrix. TPM values of known lncRNAs^104^ were extracted from the resulting matrix and averaged for each lncRNA across the replicates. Further, we binned all the expressed lncRNAs into two groups based on their median TPM value. We compared the number of POP-seq peaks (normalized per unit length of the lncRNA) mapped to the two groups of lncRNAs categorized based on low and high median expression in K562 cells. The difference in normalized peak counts between the two groups was statistically tested using Wilcoxon test^100^.

Additionally, we downloaded the raw RNA sequencing dataset of ‘whole blood’ cohort from 141 individuals from the GTEx^105^ and 174 AML patient samples from The Cancer Genome Atlas (TCGA)^81^. We processed the dataset using the NGS data processing pipeline, to generate expression levels for all human genes annotated in the human genome (hg38). We extracted the expression level of lncRNA - RP11-301G19.1 (ENSG00000227706) from the two groups; AML and GTEx ‘whole blood’. The difference in expression levels was statistically examined between the two groups using Wilcoxon test. We also calculated the patient’s survival over time using the expression levels of this lncRNA in AML patients using the Kaplan-Meier method implemented in ‘The survival’, an R package^83^. We generated a genomic track for this lncRNA using IGV and illustrated the regulatory sites identified by POP-seq. We re-investigated the genomic coordinates of this gene in SliceIt^36^ and added a panel to enable all the possible regulatory sites captured by eCLIP of RBPs in ENCODE project^28^.

## Data Availability

All POP-seq data have been deposited under Gene Expression Omnibus (GEO) accession number GSE142460. Source code of the POP-seq data processing pipeline and genome track browser shots were made available in GitHub (https://github.com/Janga-Lab/POP-seq).

## Acknowledgements

We thank the lab members for their valuable suggestions over the course of this project. We are thankful to Dr. Mark Kaplan for providing access to space and equipment to conduct this study. We also thank Swapna Vidhur Daulatabad for providing the Kaplan-Meier plot of lncRNA RP11-301G19.1.

## Authors’ contribution

MS, RS and SCJ conceived and designed the study. MS developed the POP-seq method with three versions in K562 cells and generated the NGS library for sequencing. RS implemented the bioinformatic tools and integrated the datasets for downstream data analysis. MS, RS and SCJ interpreted the data and wrote the manuscript. All authors read and approved the final manuscript.

