## Supplementary material for "Transcriptome-wide high-throughput mapping of protein-RNA occupancy profiles using POP-seq": Supplementary_materials.docx

**Supplementary Figures**

**Figure S1.** Scatter plot showing a comparison of the aligned reads per 10 kb genome between the replicates (with spearman correlation R^2^ values) for each protocol.

**Figure S2.** Comparison of POP-seq peaks with ChIP-seq and CLIP-seq data available for K562 cells (from ENCODE project). (A) Box plot showing the percentage of ChIP-seq peaks of 67 proteins and CLIP-seq peaks of 79 proteins peaks overlapped with POP-seq peaks. Level of significance between the groups were statistically tested using Wilcoxon rank sum test. (B) Enrichment analysis of POP-seq captured locations in CHIP-seq and CLIP-seq peaks of the 18 proteins than 5 random peak profiles (See methods). Significance levels (-log10 p-value) of POP-seq signals overlapping with the CLIP-seq profile and ChIP-seq profile of these 18 proteins were compared and represented as barplot for each protocol with the error bars showing the results across the random replicates.

**Figure S3.** Bar plot showing the enrichment of POP-seq peaks in eCLIP profile of individual RBP in K562 cells compared to 5 random peak profiles and statistically tested using Fisher’s exact test.

**Figure S4.** Comparative analysis of POP-seq peaks (per replicates) with the (A) lncRNA and (B) non-LncRNA gene expression. Violin plot showing the number of POP-seq peaks (normalized per unit length of the gene) binned in low and high expression groups i.e. below and above the median expression in K562 cells. The difference in normalized peak counts between the two groups was statistically tested using Wilcoxon test.

**Figure S5.** Genomic tracks showing the distribution of POP-seq variant peaks in RP11−301G19.1 along with the eCLIP profile of multiple RBPs in K562 cells.

**Figure S6.** Genomic tracks showing the distribution of POP-seq variant peaks in (A) MALAT1 and (B) NEAT1 along with the eCLIP profile of multiple RBPs in K562 cells. Total RNA-seq data (from K562 cells) is included in the track as a background control (Y-axis adjusted to POP-seq scale).

**Supplementary Table**

**Table S1.** This table summarizes the enrichment analysis of POP-seq peaks with the CLIP-seq peaks of 97 RBPs using Fisher’s exact test. For each comparison, bedtools intersect command (and enabling ‘-f ’ and ‘-r’ flags) was employed to compute the peaks (POP-seq and 5 random peak profiles, see methods) overlapped with CLIP-seq and organized as confusion matrix.

**Table S2.** This table summarizes the enrichment analysis of POP-seq peaks with the f-RIP-seq peaks of 24 RBPs using Fisher’s exact test. For each comparison, bedtools intersect command (and enabling ‘-f ’ and ‘-r’ flags) was employed to compute the peaks (POP-seq and 5 random peak profiles, see methods) overlapped with f-RIP-seq and organized as confusion matrix.

**Table S3.** This table summarizes the proportion of POP-seq peaks upon varying end-to-end peak length overlap with the ribo-seq peaks using bedtools intersect command (and enabling ‘-f ’ and ‘-r’ flags). For each comparison, peaks having length > 50 bp were filtered out from both the dataset.
