## Supplementary figures and images for "Transcriptome-wide high-throughput mapping of protein-RNA occupancy profiles using POP-seq"

### Figure S1.pdf

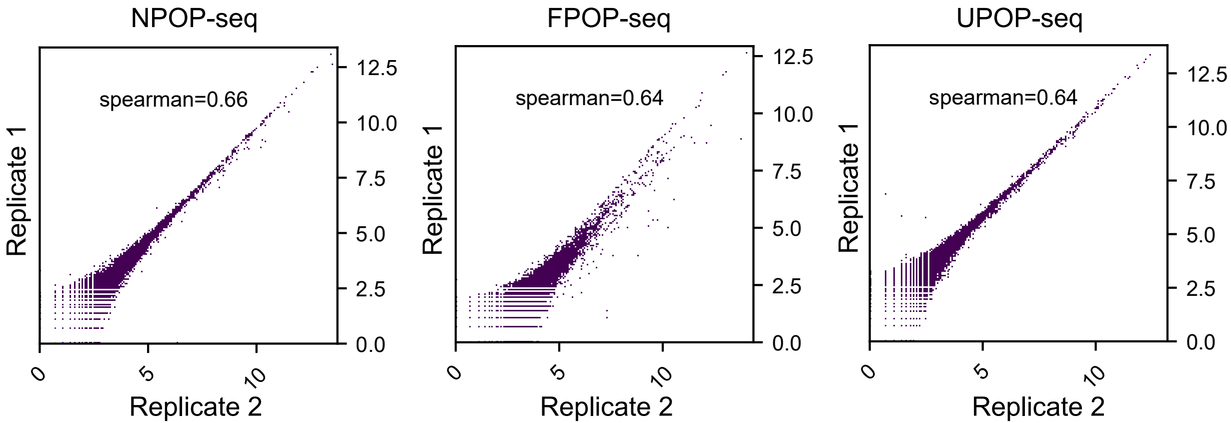

### Figure S2.pdf

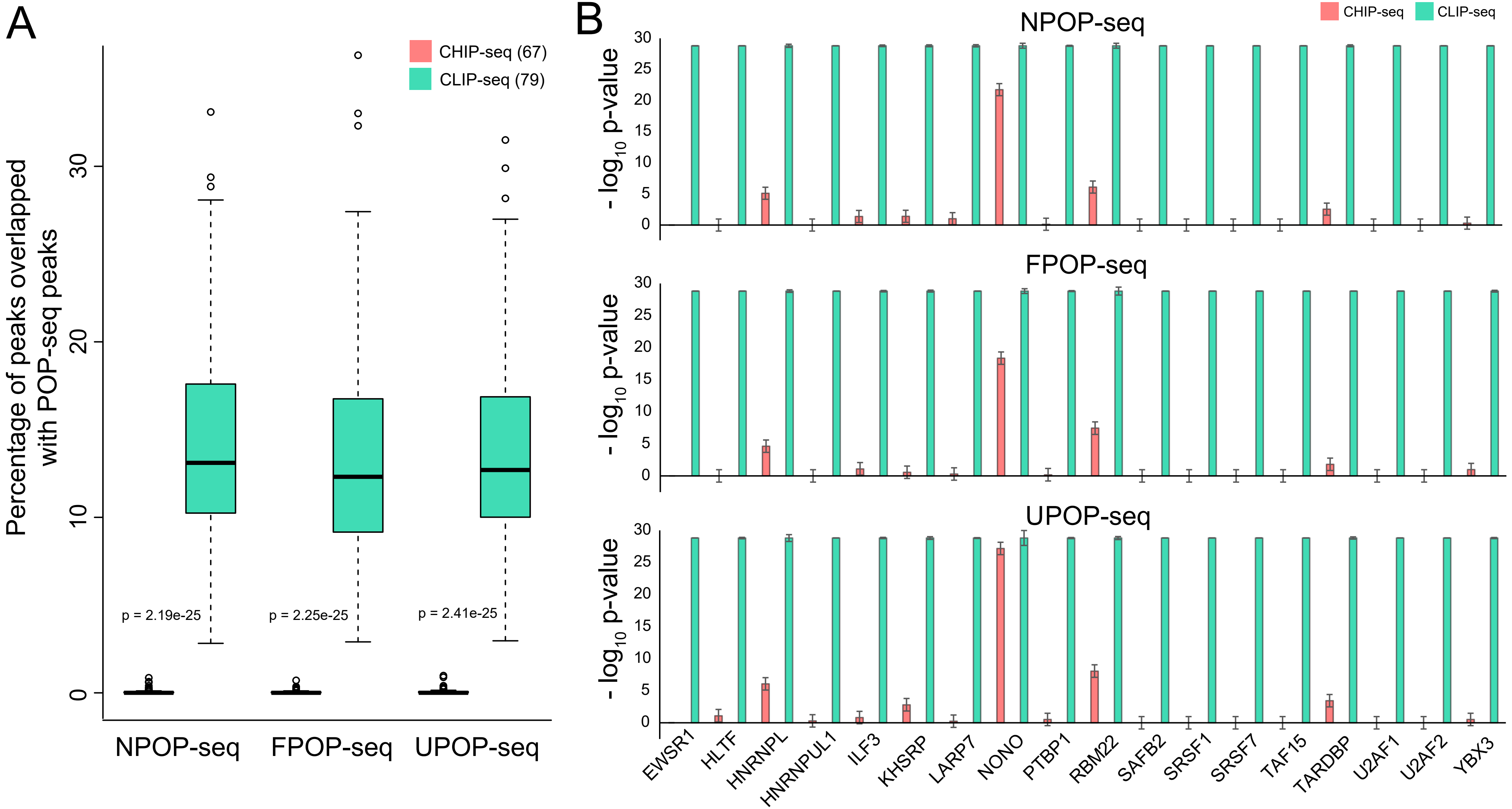

### Figure S3.pdf

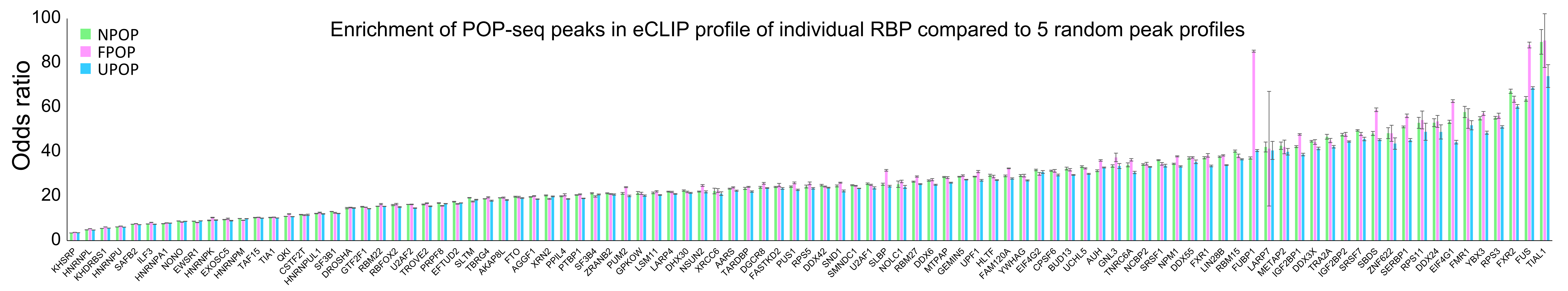

### Figure S4.pdf

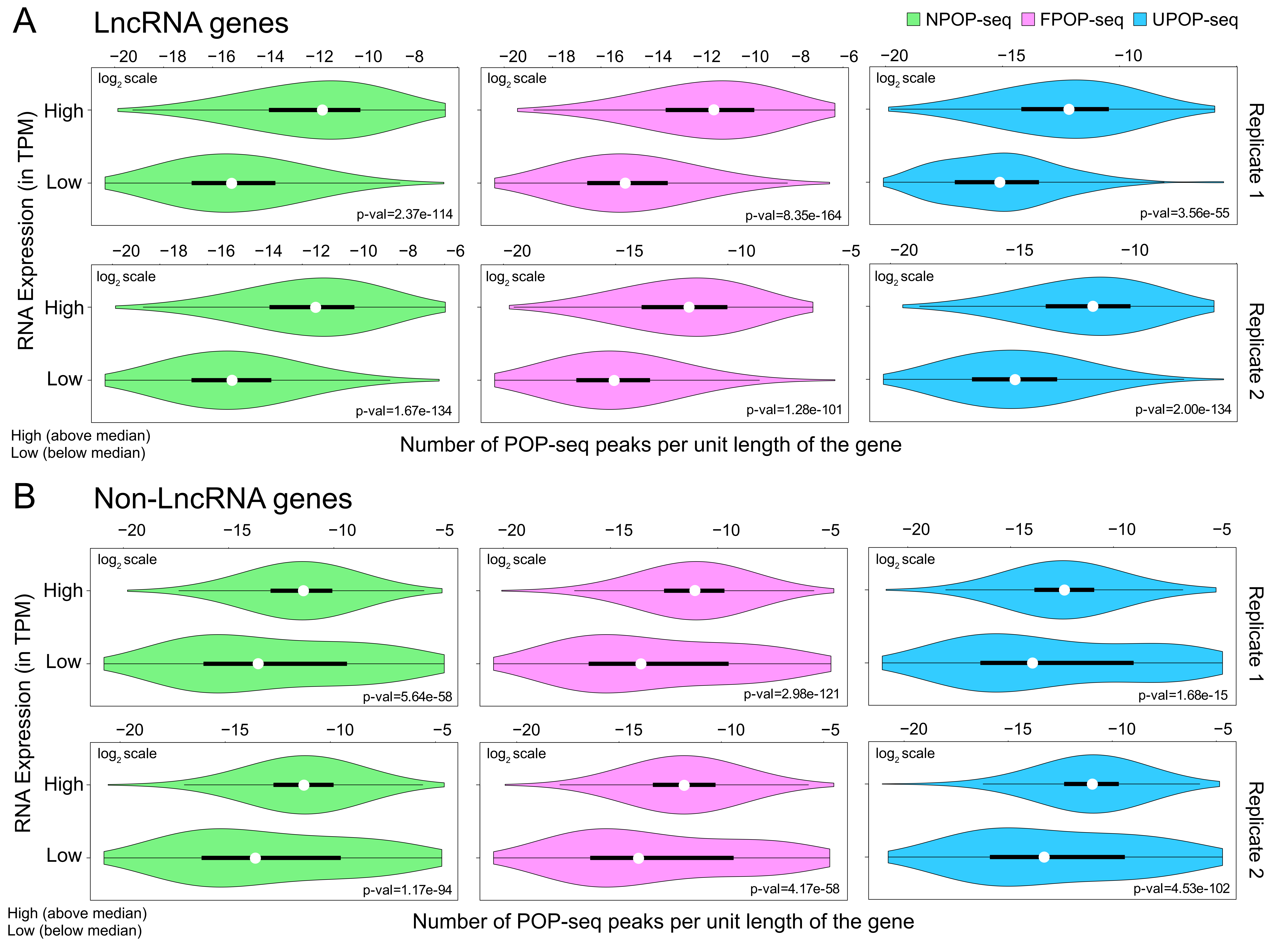

### Figure S5.pdf

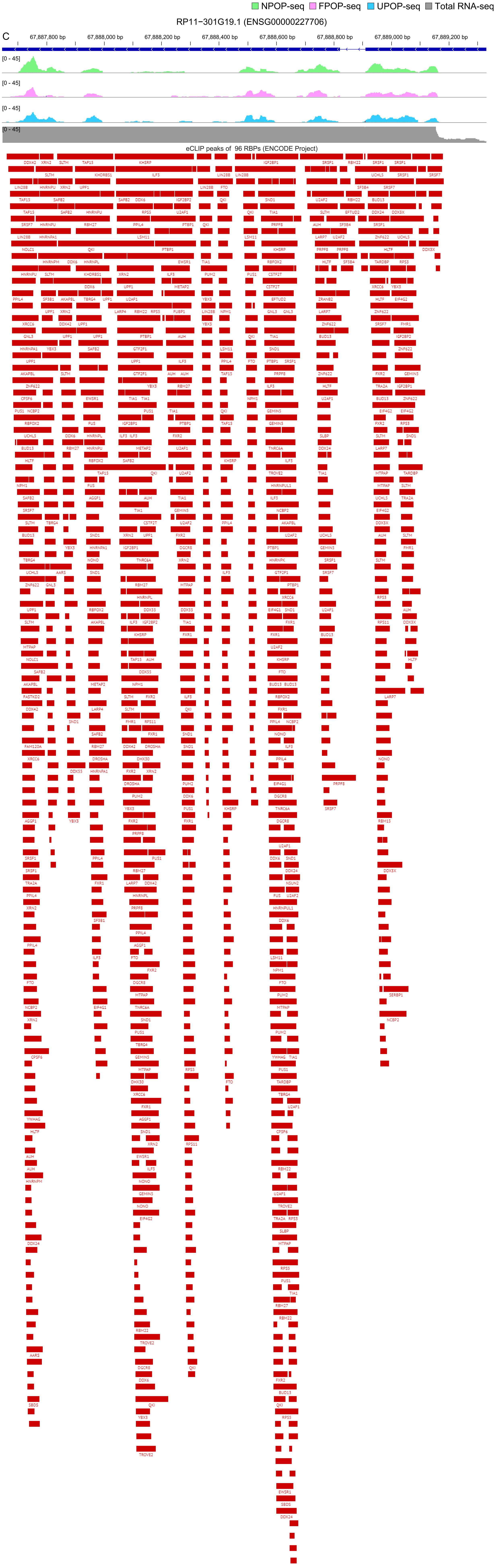

### Figure S6.pdf

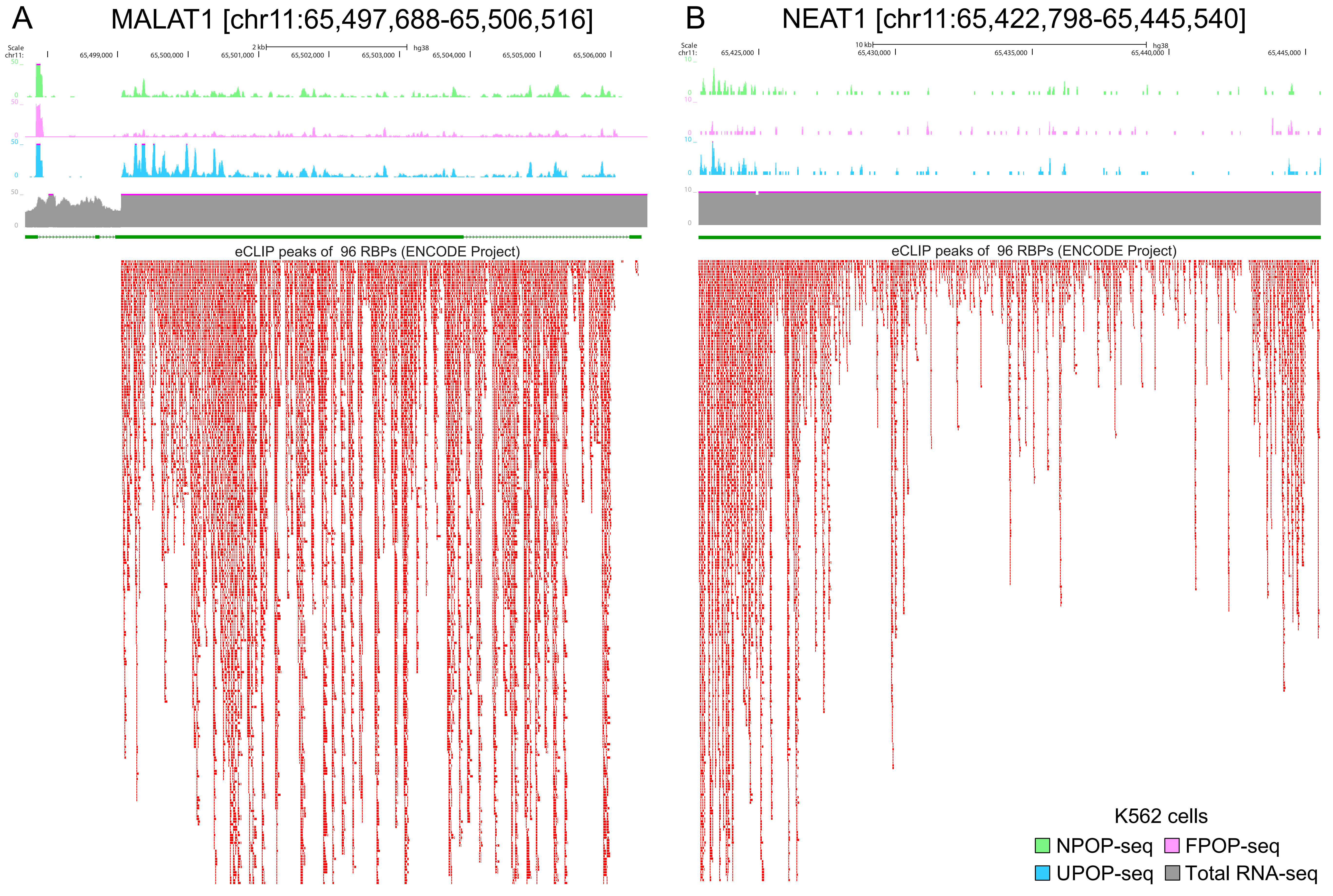
